# Variable Number Tandem Repeats (VNTRs) as modifiers of breast cancer risk in carriers of *BRCA1* 185delAG

**DOI:** 10.1101/2022.06.01.494371

**Authors:** Yuan Chun Ding, Aaron W. Adamson, Mehrdad Bakhtiari, Carmina Patrick, Jonghun Park, Yael Laitman, Jeffrey N. Weitzel, Vineet Bafna, Eitan Friedman, Susan L. Neuhausen

## Abstract

Despite substantial efforts in identifying both rare and common variants affecting disease risk, in the majority of diseases, a large proportion of unexplained genetic risk remains. We propose that variable number tandem repeats (VNTRs) may explain a proportion of the missing genetic risk. Herein, we tested whether VNTRs are causal modifiers of breast cancer risk in 347 female carriers of *BRCA1* 185delAG, an important group given their high risk of developing breast cancer. We performed targeted-capture to sequence VNTRs, called genotypes with adVNTR, and tested the association of VNTRs and breast cancer risk using Cox regression models. Of 303 VNTRs that passed quality control checks, 4 VNTRs were significantly associated with risk to develop breast cancer at false discovery rate [FDR] < 0.05 and an additional 4 VNTRs had FDR < 0.25. After determining the specific risk alleles, there was a significantly earlier age at development of breast cancer in carriers of the risk genotypes compared to those without the risk genotypes for seven of eight VNTRs. Results from this first systematic study of VNTRs demonstrate that VNTRs may explain a proportion of the unexplained genetic risk for disease and have larger effects than SNPs.

## INTRODUCTION

For carriers of pathogenic variants (PVs) in *BRCA1*, the lifetime risk for developing breast cancer (up to 80% lifetime risk) is a six-fold increase over that of average risk women and ovarian cancer risk (up to a 44% lifetime risk) is up to a 30-fold increase [1]. Despite these substantially elevated risks, penetrance is incomplete (not all carriers will develop cancer) and age at cancer diagnosis varies. The limited understanding of factors that modify cancer risks in *BRCA1* carriers hampers clinical decision-making ability, including decisions about the appropriate type and timing of risk reducing surgeries. Therefore, there is a critical, clinically relevant need for more refined risk estimates.

The variation in risk, even in identical PVs carriers, suggests that modifier factors, both genetic and environmental, affect cancer risks [2]. Studies to identify “modifier genes” that govern the phenotypic expression of *BRCA* PV carriers have been ongoing since the early 2000’s, conducted largely through the Consortium of Investigators of Modifiers of *BRCA1/2* (CIMBA) [3, 4]. Through genome-wide association studies (GWAS), single nucleotide polymorphisms (SNPs) have been identified that, when combined into a polygenic risk score (PRS), better define *BRCA1* carriers at higher and lower risk of developing breast cancer (e.g., [5-7]). However, these modifier variants explain only a portion of the variation in risk [8, 9]. Identifying additional genetic modifiers will facilitate better risk estimates for clinical decision-making on timing and options for risk reduction.

Variable number tandem repeats (VNTRs) may plausibly account for some of the missing genetic risk. They are known to modulate biologic processes, including gene expression and protein function [10-14]. These eVNTRs (VNTR expression Quantitative Trait Loci) also mediate risks of developing various cancers [15, 16] including breast cancer [17-20]. A genome-wide investigation of VNTRs as modifiers has been hampered by technical difficulties; however, adVNTR [10, 21] became available to genotype VNTRs (i.e., count repeat units) from next generation sequencing (NGS) data. This tool uses Hidden Markov models (HMM) to model each VNTR, count repeat units, and detect sequence variation.

We tested a new paradigm – that VNTRs are causal modifiers of breast cancer risk. They have not been systematically investigated as they are poorly tagged by nearby SNPs [12]. Previous GWAS conducted through CIMBA have demonstrated heterogeneity of breast cancer risk by *BRCA1/2* mutations, breast tumor subtypes, and race and ethnicity [22]. Therefore, to reduce potential confounding with unmeasured variables, we tested the association in carriers of a single recurring PV in *BRCA1*. We performed targeted-capture to sequence VNTRs, called genotypes with adVNTR, and explored the association of VNTRs and breast cancer in in 327 women carrying the pathogenic *BRCA1* 185delAG mutation NM_007294.4(*BRCA1*):c.68_69del (p.Glu23fs) (rs80357914).

## METHODS

### Cases

Females carrying the pathogenic *BRCA1* variant 185delAG (NM_007294.3:c.66_67del) were eligible. Of the 347 participants with DNA, 250 were enrolled by Dr. E. Friedman from the Suzanne Levy-Gertner Oncogenetics Unit at the Sheba Medical Center (SMC) in Israel. All participants underwent oncogenetic counseling and genotyping of cancer susceptibility genes, including *BRCA1*. Referral to the oncogenetics services came from several sources: women who developed breast and/or ovarian cancer (consecutive women at the SMC), cancer free women with a significant family history of breast and /or ovarian cancer, and from population screens of the three predominant mutations in Ashkenazi Jewish (AJ) women in *BRCA1* and *BRCA2* [23], a procedure recently approved and included in the Israeli “health basket” for all AJ women as a screening procedure with no need for pre-test counseling. Another 95 participants were recruited and enrolled by Dr. J. Weitzel into the Clinical Cancer Genomics Community Research Network housed at the City of Hope and another 2 participants were recruited and enrolled in a research study led by Dr. S. Neuhausen. All participants provided written informed consent under IRB-approved protocols at their respective institutions. All participants were unrelated.

#### VNTR genotyping

##### VNTR selection

To get an initial list of VNTRs (of four or more base pair repeats), Tandem Repeat Finder (TRF) [24] was applied to the human reference genome [GRCh38], and 559,804 VNTRs were identified. To focus on the most relevant candidates, we selected VNTRs that intersected with coding exons, promoters, or untranslated regions (UTRs) of genes in RefSeq (https://www.ncbi.nlm.nih.gov/refseq/).VNTRs were excluded if they were located in low-complexity sequence (e.g. close to a telomere) resulting in 8953 candidate VNTRs. Lastly, only candidate VNTRs with total length of 140 bp or shorter (n=6271) were included so that genotypes could be confidently assigned with Illumina short read sequencing data. We used the Agilent SureDesign software to design probes for 6271 VNTRs. Of these 6271 VNTRs, 1398 are in coding exons, 2000 are in promoter regions, and 2873 are in UTRs. Using the least stringent parameters to design probes for target enrichment, probes were designed to cover 6186 VNTRs; 85 VNTRs were excluded because they were in repetitive DNA regions where probes could not be designed. We also excluded 21 VNTRs on the Y chromosome.

##### Library preparation

Libraries were created from 500 ng DNA using KAPA Hyper (KAPA Biosystems) reagents along with our optimized protocols [25, 26] to maximize efficiency and minimize cost including bar-coding samples prior to hybridization in order to hybridize 24 samples per bait capture kit. Prior to bait capture, indexed samples were carefully quantified with both Picogreen and qPCR assays to ensure equal representation of each sample in the pool. The protocol has been optimized to minimize the number of PCR cycles, reducing duplicate reads to less than 20%. Sequencing was performed in the City of Hope Integrative Genomics Core (IGC) on a HiSeq2500 genetic analyzer (Illumina Inc, San Diego, CA).

##### Targeted capture DNA sequencing and processing of reads

For the first 192 samples, we obtained sequencing on 150 bp paired-end reads. The remaining samples were sequenced with 250 bp paired-end reads in order to provide more flanking sequence for better accuracy of the call.

Once sequencing was complete, the fastq files were uploaded to a shared folder for processing. Sequence reads were aligned to NCBI build GRCh38 using Burrows-Wheeler Aligner (BWA) to generate BAM files. Duplicate reads were detected by Picard MarkDuplicates (http://broadinstitute.github.io/picard/) and only unique reads were kept for subsequent analysis. The BAM files including unmapped reads were used for assigning VNTRs. From the BAM files, genotypes from VNTRs were assigned using adVNTR-NN adapted from adVNTR [21] based on minimal total supporting reads ≥ 10 and minimal proportion of reads to support alternative allele ≥ 0.25.

##### Confirmation of VNTR genotyping results from adVNTR

Using the unique flanking regions of the selected VNTRs, PCR primers were designed to amplify 50 ng DNA from up to 4 samples per VNTR genotype. PCR reactions were performed using TAQ polymerase (Qiagen) and amplification was confirmed using gel electrophoresis. Samples were then sequenced on an Applied Biosystems SeqStudio Genetic Analyzer (ThermoFisher Scientific).

VNTR sequences were visualized using Quality Check and Variant Analysis Modules on the ThermoFisher Cloud. The visualized sequence in conjunction with the product sizes from the post-PCR gel electrophoresis were used to verify genotyping calling made by adVNTR. For homozygotes, this was done by observing a single band of the correct size during gel electrophoresis and by quality sequence for the number of repeats called by adVNTR. Whereas heterozygotes were confirmed by observing multiple bands of expected size differentials on the gel and a poor-quality sanger sequence at the point of allele differences.

##### Statistical analysis

After genotypes were assigned for each VNTR, we tested for HWE. For those that were in HWE (p > 0.001), we tested the association of the VNTR and risk to develop breast cancer using Cox regression models in both the primary and secondary analyses (described below). In the model, women with a first breast cancer are considered as affected with time to breast cancer diagnosis as the end point; those unaffected with any cancer or diagnosed with ovarian cancer prior to breast cancer were censored at age of last follow-up and age at ovarian cancer diagnosis, respectively. There were too few cases of ovarian cancer for association analysis.

In the primary analysis, we tested the association between each VNTR marker as a continuous variable and disease risk. Three separate VNTR genotypes were constructed: 1) the average length of the two alleles; 2) the length of only the shorter allele; and 3) the length of only the longer allele [27]. Analyses were adjusted for sample collection site (US or Israel). Probability values were adjusted for multiple comparisons using the False Discovery Rate (FDR) method of Benjamini and Hochberg [28]. For VNTRs with associations with FDR < 0.25 in the primary association analysis, a secondary association analysis was performed to identify the specific risk groups of repeat alleles using a sliding window method [27]. Specifically, a threshold T along the number of repeats from short to long was used to dichotomize allele lengths. An allele was denoted as ‘short’ if it had shorter than T repeat motifs, and ‘long’ otherwise. Multiple values of threshold T were chosen for association tests. For each specific threshold T, the VNTR genotype of an individual was converted to homozygous-short-allele genotype (S/S), heterozygous-short-and-long-allele genotype (S/L), or homozygous-long-allele genotype (L/L). The optimal threshold for each VNTR was determined by choosing T that provided the smallest p-value among the multiple association tests. This secondary analysis allowed us to identify critical cut points for risk alleles along the continuous repeat allele distribution in a VNTR and then to estimate the effect size of the association related to the risk genotypes. Kaplan-Meier curves and log-rank tests were used to graphically examine differences in the cumulative probability of breast cancer risk among VNTR genotype groups categorized using the critical cut points for risk alleles.

##### Luciferase assays

We conducted luciferase assays to test alleles of one VNTR in the promoter region to determine if it affected expression. We selected the VNTR with the lowest FDR that was in a promoter or 5’UTR region.

##### PCR amplification and subcloning

Primers, designed to flank the VNTR, also included restriction enzyme sites. The PCR products for each risk allele and reference repeat allele served as inserts for subcloning into plasmids. PCR reactions were performed using Herculase high fidelity polymerase (Agilent) and the products sizes were confirmed by gel electrophoresis. Because an individual with a rare VNTR allele is almost always heterozygote, the PCR reaction generated two different sized products. Following amplification, each PCR product was cloned separately into the pGEM-T plasmid (Promega) according to the manufacturer’s instructions followed by transformation into *E. coli* cells. Ampicillin resistant colonies were picked, grown overnight, followed by the isolation of plasmid DNA. Several plasmids were Sanger sequenced to ensure that clones contained inserts with the appropriate sequence and length corresponding to the desired repeat alleles.

##### Construction of luciferase reporter plasmids

Following sequence confirmation, the pGEM-T plasmids served as the template DNA for another PCR reaction using the same primers described above. The PCR products containing the VNTR alleles were digested with the appropriate restriction enzymes, purified, and cloned into restriction enzyme sites in the pGL3-Promoter luciferase gene-reporter plasmid (Promega) upstream of the SV40 promoter. Sequencing was performed to confirm the fidelity and orientation of all inserts.

##### Luciferase assay

We transfected the luciferase gene-reporter plasmids along with the pRL Renilla luciferase control plasmid into an MCF7 cell line (from ATCC) and performed luciferase assays using the Dual-Luciferase Reporter Assay System (Promega). After 48 hours, each group of cells was lysed and manipulated according to the manufacturer’s instructions, and the fluorescence intensity of each group of cells was expressed as the ratio of firefly luciferase activity to Renilla luciferase activity. The relative luciferase activities of the plasmid constructs were determined by normalizing the standardized values to empty pGL3-Promoter plasmid. All transfections were performed in quadruplicate, and each construct was tested in three independent experiments. The average of the 12 relative luciferase measurements for each allele were expressed as the mean ±standard error of mean (SEM). Difference in relative activity values between the risk repeat allele group and reference repeat allele group was tested by one-way ANOVA analysis. The P-value was adjusted for multiple testing using the Tukey’s method [29]; adjusted p-values less than 0.05 were considered as statistically significant.

## RESULTS

### Participants

The cancer status and ages at diagnosis or enrollment (for non-cancer cases) are shown in Table 1. Of the 347 women, ages ranged from 18 to 77 years with 49% having been diagnosed with breast cancer, of which 4.9% also were also diagnosed with ovarian cancer. The median age at diagnosis was 55 years (95% CI: 50 - 58 years). The cumulative risk of breast cancer by age 70 years was 75%.

**Table 1.** Participant characteristics

|  | N (%) | Age in years (Mean, Range) |
| --- | --- | --- |
| Sample origin |  |  |
| US | 97 (28.0) | 43.9, 18 - 72 |
| Israel | 250 (72.0) | 46.8, 24 - 77 |
| Cancer status |  |  |
| Breast | 153 (44.1) | 43.5, 24 - 75 |
| Ovarian | 60 (17.3) | 54.0, 35 - 77 |
| Breast & Ovarian | 17 (4.9) | 48.0, 27 - 68 |
| Unaffected | 117 (33.7) | 44.9, 18 - 76 |

### VNTR genotyping

In total, we sequenced 6165 VNTRs in 347 *BRCA1* 185delAG PV carriers. Genotypes were called using adVNTR-NN. In Figure 1, the flow diagram of steps for elimination of VNTRs and samples is shown. Of 6165 VNTRs, 3847 (62.4%) VNTRs were removed due to missing more than 5% of genotypes, with the main reasons being VNTRs located in GC-rich regions which had poorer amplification during library generation, imperfect repeats, or flanked by other repetitive elements. Another 1622 VNTRs were removed because they were monomorphic (1588 VNTRs) or not in Hardy-Weinberg equilibrium (P value < 0.001; 34 VNTRs). Lastly 393 VNTRs had heterozygosity < 0.02. Because this is a homogeneous dataset of Ashkenazi Jewish ancestry, it was expected that more VNTRs would be monomorphic and within VNTRs, not all alleles would be present. Twenty samples were removed that had more than 10% missing genotypes leaving 327 for analysis. The summary of repeat alleles in this dataset for the 303 VNTRs is shown in Table 2.

**Figure 1.**
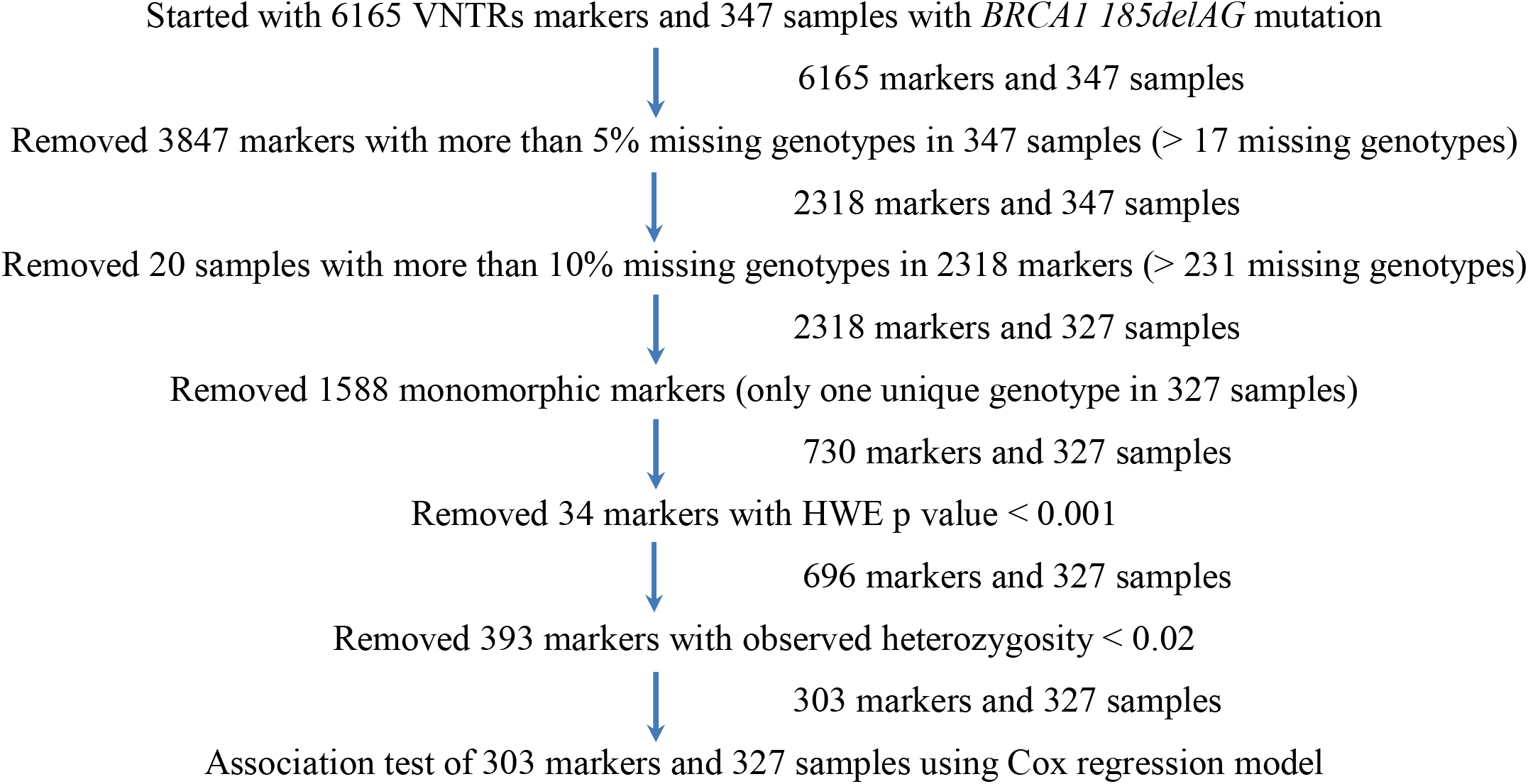
Flow diagram of process and result of VNTR marker and sample filtering.

**Table 2.**
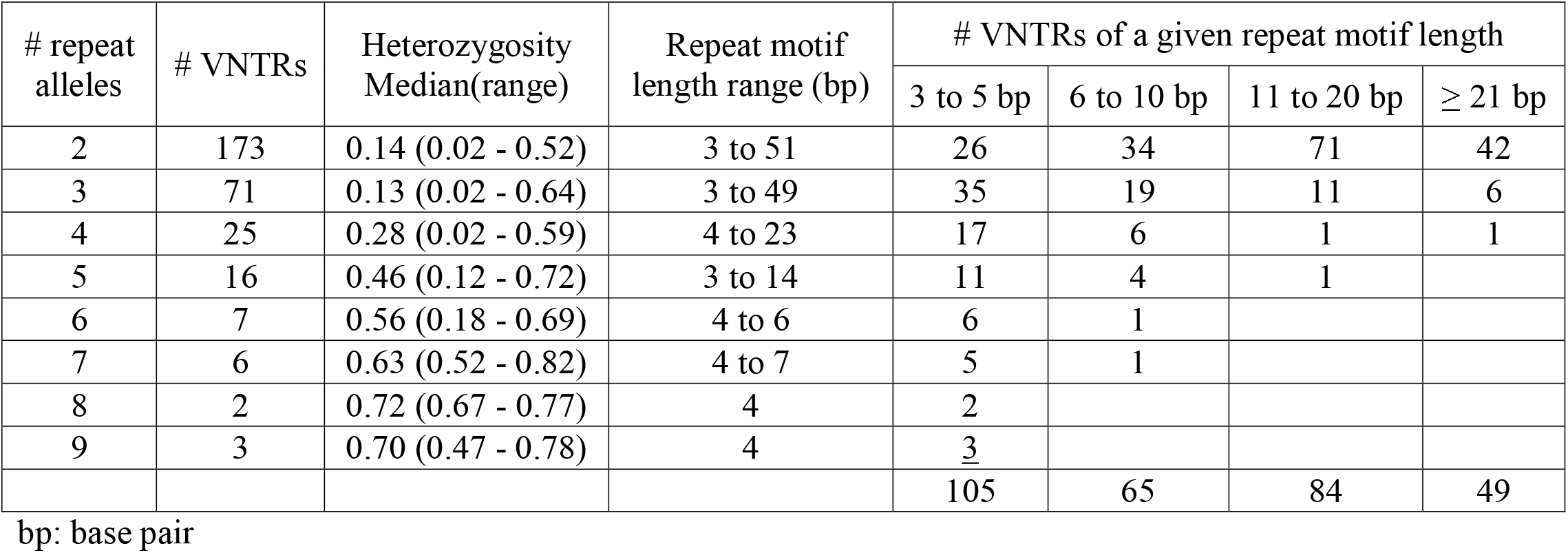
Summary of repeat alleles in the 303 VNTRs in 327 female *BRCA1 185delAG* mutation carriers

### Association of VNTRs and risk of developing cancer

In the primary analysis, we used Cox proportional hazards models to evaluate the association between each VNTR and risk of developing breast cancer, considering the VNTR as a continuous variable. Of 303 VNTRs analyzed, 8 VNTRs had unadjusted p-values < 0.005, of which four VNTRs had FDR < 0.05, and an additional four had FDR < 0.25 (Table 3; Supplemental Table 1). The alleles for each of the eight VNTRs were accurately called, with 100% consistency among the adVNTR, agarose gel, and Sanger sequencing results (VNTR 558420 is shown as an example in Supplemental Figure 1). We then conducted the secondary analysis for the eight VNTRs to identify the specific risk repeat alleles contributing to the significant association. For 7 of 8 VNTRs, there was a significant (P < 0.05) difference in breast cancer risk between risk and reference genotype groups categorized using the critical cut point of risk allele (forward slash character in Table 3). For VNTR 47260, although breast cancer risk increased with repeat length based on the linear trend test (FDR = 0.035), there were too few long repeat alleles (> 9R) for a stable estimation of cut point in the categorical test. Kaplan-Meier (KM) curves were used to graphically show the difference in the cumulative probability of breast cancer risk for the VNTR genotype groups (Figure 2 and Supplemental Figure 2). Individuals with the risk genotypes had significantly earlier ages at developing breast cancer (log-rank p value < 0.05) (Figure 2 and Supplemental Figure 2). For example, the median ages at breast cancer diagnosis for carriers with the S/S genotype and the L/L genotype in VNTR357331 were 40 years and 56 years, respectively (log-rank p value of 0.0014, Figure 2), indicating the age-modifying effect of breast cancer diagnosis among carriers harboring risk genotype (S/S).

**Table 3.** Association of VNTR with breast cancer risk in female carriers of *BRCA1 185delAG*

| VNTR ID<sup>a</sup> | Trend test, VNTR as a continuous variable |  |  | Categorical test to determine cut point of risk repeats in a VNTR |  |  |  |
| --- | --- | --- | --- | --- | --- | --- | --- |
|  | P | FDR | HR (95% CI) | Repeat number / critical cut point<sup>b</sup> | Reference, risk genotype<sup>c</sup> | P | HR (CI) |
| 253688 | 9.1E-05 | 0.024 | 6.12 (2.47 - 15.15) | 5, 9 / 10 | S/S, S/L | 9.1E-05 | 6.12 (2.47 - 15.15) |
| 357331 | 2.6E-04 | 0.035 | 3.57 (1.79 - 7.14) | 4 / 5, 6 | L/L, S/S | 2.9E-04 | 3.59 (1.80 - 7.16) |
| 472060 | 3.9E-04 | 0.035 | 2.54 (1.52 - 4.25) | 4 / 9, 13 | L/L, S/L | 6.8E-01 | 0.81 (0.30 - 2.19) |
| 412033 | 6.9E-04 | 0.046 | 1.35 (1.14 - 1.59) | 7, 8 / 9, 10, 11 | L/L, S/S | 3.4E-02 | 1.69 (1.04 - 2.75) |
| 558420 | 2.5E-03 | 0.135 | 6.06 (1.88 - 19.51) | 2, 3 / 4 | S/S, S/L | 2.5E-03 | 6.06 (1.88 - 19.51) |
| 735300 | 4.1E-03 | 0.183 | 3.08 (1.43 - 6.65) | 5 / 6 | S/S, S/L | 4.1E-03 | 3.08 (1.43 - 6.65) |
| 945060 | 4.8E-03 | 0.183 | 1.60 (1.15 - 2.23) | 6, 7, 8, 9 / 10 | S/S, S/L | 1.1E-02 | 2.73 (1.26 - 5.94) |
| 549198 | 2.0E-03 | 0.209 | 2.94 (1.49 - 5.92) | 4 / 5 | L/L, S/L | 2.0E-03 | 2.97 (1.49 - 5.91) |

**a.** VNTR IDs were assigned in this project and their genomic annotations are provided in supplemental table 1; **b:** in categorical test to determine cut point of risk repeat for a VNTR by a sliding window method, the forward slash “/” character is the final critical cut point based on smaller p values among the multiple tests of cut points, greater or equal to repeat number at right side of slash is defined as long allele (L), repeat numbers at left side of slash as short allele (S); **c:** in Cox regression model using categorical VNTR genotype as predict variable, hazard ratio was calculated by comparing risk genotype to reference genotype.

**Figure 2.**
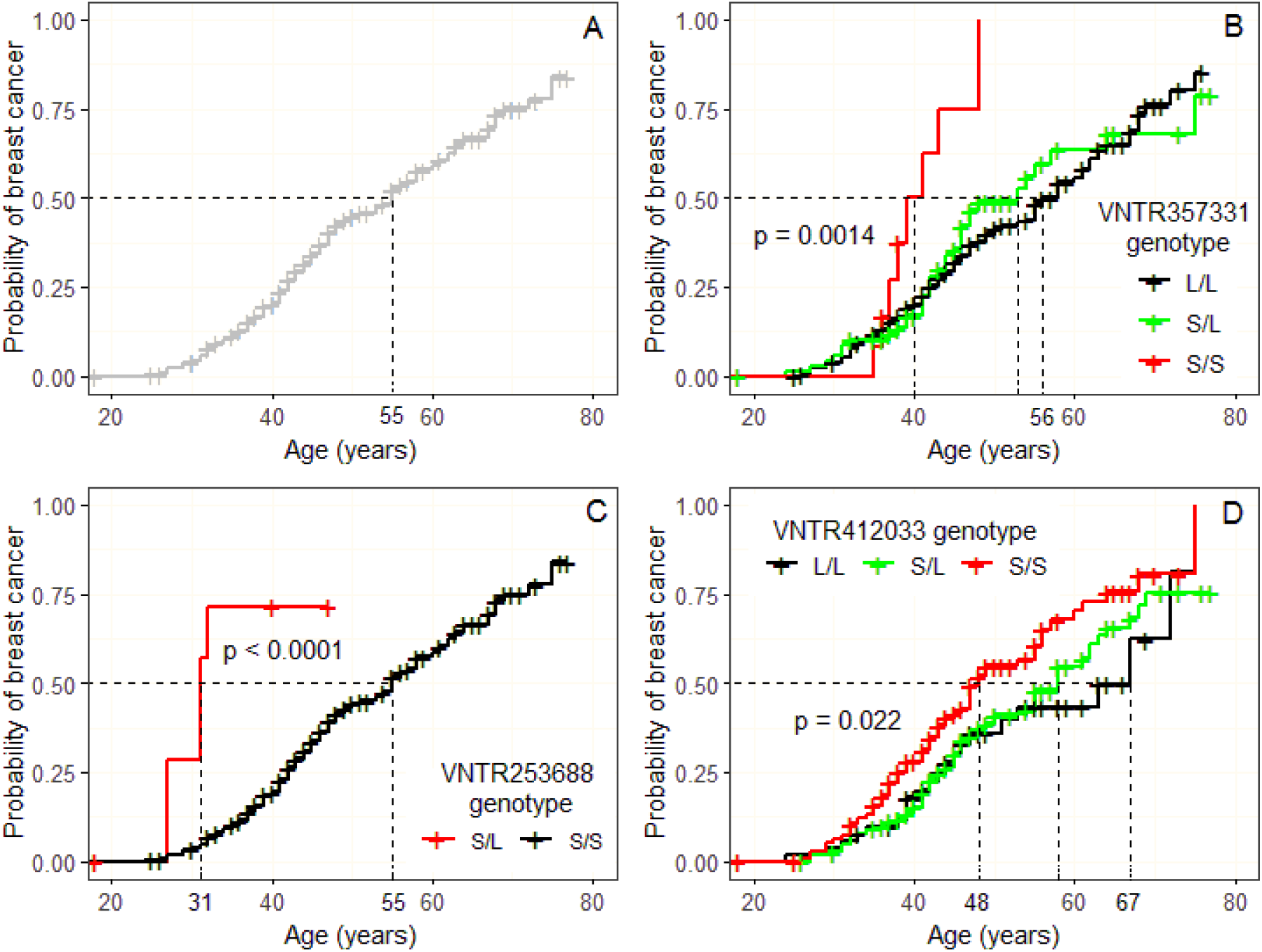
Kaplan-Meier estimates of the cumulative probability of breast cancer diagnosis. The age at breast cancer diagnosis is on the X-axis and proportion of participants diagnosed with breast cancer is on the Y-axis. The horizontal**/**vertical dash line is the median age at diagnosis of breast cancer. In this step function of breast cancer risk over age, in panel A, the cumulative incidence of breast cancer for all *BRCA1* 185 delAG carriers is shown. For panels B, C, and D, the Kaplan-Meier curves for each of the three VNTRs with FDR < 0.05 are shown. Panel B is VNTR 357331, Panel C is VNTR253688, and panel D is VNTR412033. The black line is the reference genotype, the green line (when present) is the heterozygote genotype, and the red is the risk genotype. For each of the VNTRs, there were significantly different risks by genotype (log-rank p value < 0.05) with earlier ages of developing breast cancer among participants carrying the risk genotypes.

### Effect of VNTR alleles on expression

For testing the effect on gene expression, we selected the VNTR with the lowest FDR that was located in a gene promoter or 5’ UTR. We tested VNTR 558420 located in the 5’ UTR of *ZNF501* (p-value = 0.0025 and FDR = 0.135) (Table 3 and Supplemental Figure 3) with repeats of 2R, 3R and 4R and 3 genotypes (3 samples with genotype 2/3, 314 with 3/3, and 5 with 3/4). In Figure 3, normalized luciferase activity is shown for the 2R, 3R, 4R repeats and the control (empty vector) with standard error bars on the top of each group mean. There was a significant (adjusted p value < 0.05) difference between the 2R and 4R groups with the 3R intermediate (Figure 3) and a significant linear trend of decreased luciferase activity with increasing number of repeats (p = 0.021) (Supplemental Figure 4).

**Figure 3.**
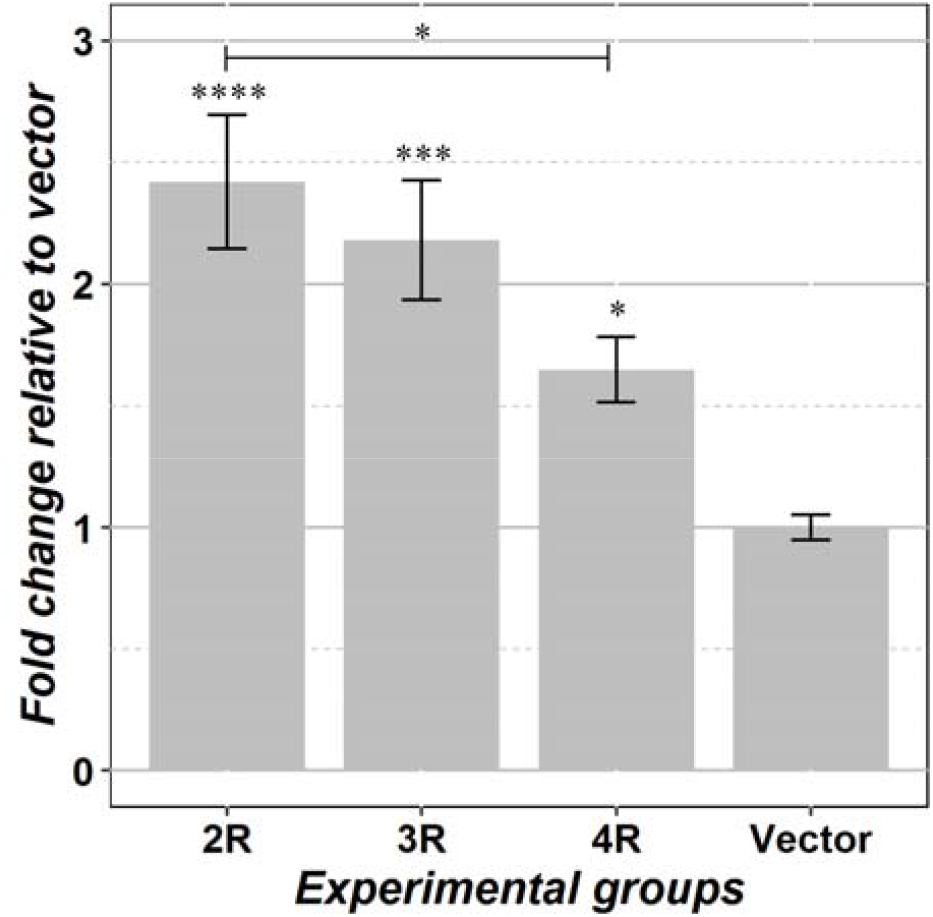
Association of VNTR558420 with *ZNF501* gene expression by luciferase assay. Each experimental group is composed of 12 data points. Data represent fold change in the repeat group relative to vector group, with standard error bar shown for each group. Significance was assessed by one-way ANOVA with pairwise t test and P-value adjusted by the Tukey’s method. Asterisk above standard error bar indicates significance test between the repeat group and vector group; asterisk above the line indicates the significance between the 2R and 4R repeat groups; *P < 0.05, ** P < 0.01, *** P < 0.001, **** P < 0.0001.

## DISCUSSION

Our study is the first to conduct a systematic study of VNTRs and association with risk to develop cancer in high-risk *BRCA1* PV carriers. We identified four VNTRs significantly associated with risk of developing breast cancer in women carrying the 185delAG *BRCA1* PV (FDR < 0.05) and another four VNTRs associated with FDR < 0.25.

None of the small number of previous association studies of risk of developing breast cancer and VNTRs at candidate genes had investigated the eight VNTRs we identified. Krontiris and coworkers reported an association of rare alleles in a *HRAS1* VNTR and development of cancers, including breast cancer [17], and a meta-analysis of 13 breast cancer studies found an association with breast cancer risk [30]. Functional analysis showed that this *HRAS* VNTR altered CpG DNA methylation [31]. In a meta-analysis of 17 studies of a CAG-repeat polymorphism in the androgen receptor, they found an association of longer CAG repeats with an increased risk of breast cancer in Caucasian women [32]. In a meta-analysis of two studies of the MNS16A VNTR in the *hTERT* promoter, they found a significant association with development of breast cancer. In a Japanese study of an 18-bp VNTR in the promoter of *PTTG1IP*, they found a signficant association with risk of estrogen-receptor positive breast cancer, with functional analysis showing that an increase in the number of repeats increased the binding affinity of ER-alpha [20]. In a study of a VNTR in the promoter of *XRCC5*, they found a significant association with age at breast cancer diagnosis [18]. Our study did not include the two trinucleotide repeats and the *XRCC5* VNTR, and the *MSN1* VNTR was monomorphic and the *PTTG1IP* VNTR was missing too many genotypes in our set and thus were excluded.

Of the eight VNTRs that we found to be associated with risk of developing breast cancer in this population, several warrant further investigation. VNTR 945060 is in the 5’UTR of *ERCC6L*, a DNA helicase. ERCC6L is highly expressed in breast tissue and higher levels of expression have been associated with worse survival [33]; silencing of ERCC6L in breast cell lines significantly inhibited cell proliferation [33, 34]. A second VNTR, 253688, is located 3’ of *FLJ22447*, a lncRNA located near *HIF-1α*. In a study of esophageal squamous cell carcinoma and gastric cancers to determine the effect of *FLJ22447* on *HIF-1α*, they observed that low expression of lncRNA was associated with expression of *HIF-1α* suggesting that *FLJ22447* may have a regulatory function on *HIF-1α* expression [35]. High over-expression of *HIF-1α* is common in breast cancers and is particularly common in *BRCA1* carriers [36-38]. This VNTR may alter risk to develop breast through affecting *HIF-1α*.

Given the reports that there are shared genetic contributions between breast cancer and schizophrenia [39], it is interesting that three of the VNTRs are at or in genes (*SYN2, ZNF501, ZNF804A*) associated with risk to develop schizophrenia [40-44]; VNTR 549198 is in exon 12 of *SYN2*; VNTR 472060 is in exon 4 of *ZNF804A*; and VNTR558420 is in the 5’UTR of *ZNF501* and all are most commonly expressed in brain (proteinatlas.org). From our luciferase assays, there was differential expression from varying alleles in the VNTR in the 5’UTR of *ZNF501*; expression differences for this VNTR were only associated with brain tissue in GTEX [10]. The exonic VNTRs in *SYN2* and in *ZNF804A* cause expansions of poly-serine (Supplemental Figure 5) and poly-alanine (Supplemental Figure 6) tracts, respectively. VNTR expansions in gene coding regions have been associated with multiple diseases [45]. Further investigation is needed to assess possible roles in development of breast cancer.

This was a pilot study to determine the feasibility of conducting targeted sequencing of VNTRs and investigating the association of VNTRs as modifiers of disease risk, similar to what has been accomplished with SNPs [8, 46]. We purposefully included women carrying the specific *BRCA1* 185delAG Ashkenazi Jewish founder PV to try to explain the known variation in risk in women carrying this PV and to reduce potential confounding with unmeasured variables; however, the consequence is that it reduced the number of VNTRs that were polymorphic and restricted the sample size. In hindsight, using targeted capture and sequencing of 250 bp reads limited the size of repeats and reduced the number of VNTRs that made it through all the quality control checks due to poor amplification of VNTRs in GC-rich regions, difficulty in aligning VNTRs with imperfect repeats and/or with low complexity/repetitive sequence in the flanking regions. As costs continue to decrease for whole-genome sequencing and for long-read sequences such as performed by PacBio, we will be able to obtain analyzable data on a larger number of VNTRs.

*BRCA1* breast cancers are generally basal, triple-negative hormone receptor cancers (TNBC). We have seen from SNP studies of both *BRCA1* carriers and women with TNBC that there are fewer SNPs associated with risk than for estrogen-receptor positive breast cancers. Thus, identification of VNTRs significantly associated with risk of developing breast cancer in this genetically and ethnically homogeneous population is encouraging; several of which have been observed to play a role in breast cancer. HRs for these VNTRs ranged from 1.7 to 6.1 whereas HRs for SNPs generally range from 1.01 to 1.4 [8, 47], suggesting that VNTRs may have larger effects than SNPs. These results need to be validated in larger datasets that include women of diverse ethnicities, a wider spectrum of *BRCA1* PVs, and carriers of *BRCA2* PVs. Moreover, a larger genome-wide VNTR association study may identify additional VNTRs.

In summary, the results from this study demonstrate that VNTRs may explain a proportion of the unexplained genetic risk for disease. Similar to SNPs, VNTRs significantly associated with the disease of interest could be incorporated into polygenic risk scores (PRS) to test for improved risk assessment and clinical applicability.

Acknowledgments

This work was supported by the Basser Foundation (SLN and EF) and the Beckman Research Institute of City of Hope Excellence Awards Program. Sequencing of the VNTRs was performed in the Integrative Genomics Core supported by the National Cancer Institute of the National Institutes of Health under grant number P30CA033572. The content is solely the responsibility of the authors and does not necessarily represent the official views of the National Institutes of Health. SLN is partially supported by the Morris and Horowitz Families Professorship. VB and JP were supported in part by R01GM114362, R01HG010149 and RO1HG011558 from the NIH.

## Author Contribution Statements

SLN and EF conceived of the idea, developed the design, and obtained funding to conduct the study. VB, MB, and JP did the analysis of the sequencing data to obtain genotypes. AA and CP prepared the DNA libraries for sequencing, conducted the luciferase assays, and the validation of the VNTR genotypes. YCD did quality control of the VNTR genotypes and conducted the association statistical analysis. SLN, EF, JW, and YL contributed DNA samples and data. All authors have contributed to, read, and approved the manuscript.

