## Supplemental Figure 1 for "Variable Number Tandem Repeats (VNTRs) as modifiers of breast cancer risk in carriers of *BRCA1* 185delAG"

4

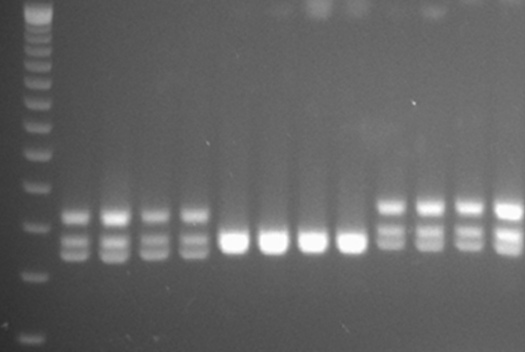

A

3

2

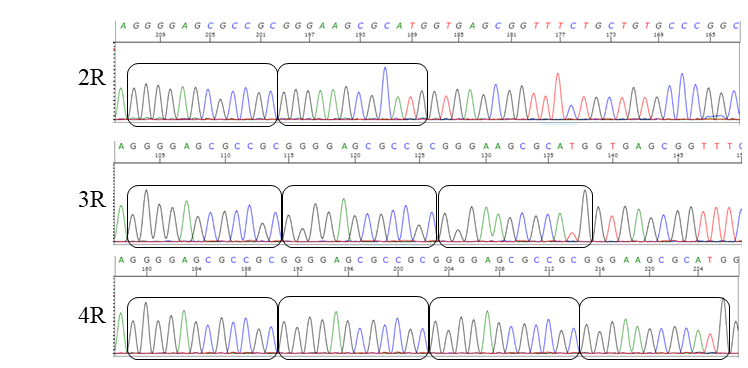

B

**Supplemental Figure 1: Confirmation of VNTR genotyping results for VNTR 558420 located in the 5’ UTR of ZNF501.** Panel A) Agarose gel image of PCR products displaying the 2R/3R, 3R/3R, and 3R/4R genotypes. The presence of the upper bands in the 2R/3R and 3R/4R genotypes is due to partial annealing of the different sized PCR products. Panel B) Sanger sequencing data displaying the 2R, 3R, and 4R repeat alleles.
