## Supplemental Figure 2 for "Variable Number Tandem Repeats (VNTRs) as modifiers of breast cancer risk in carriers of *BRCA1* 185delAG"

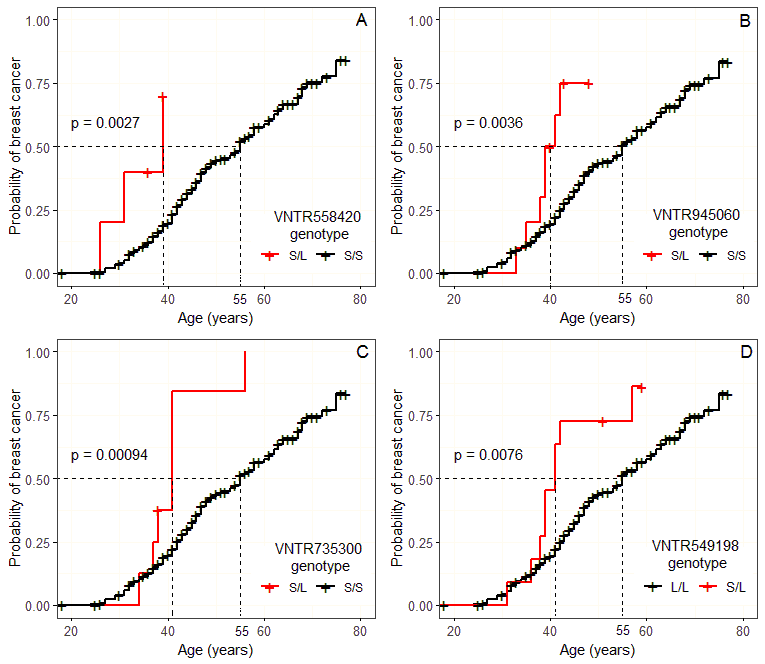

**Supplemental Figure 2. Kaplan-Meier estimates of the cumulative probability of breast cancer diagnosis.** The age at breast cancer diagnosis is on the X-axis and proportion of participants diagnosed with breast cancer is on the Y-axis. The horizontal**/**vertical dash line is the median age at diagnosis of breast cancer. In this step function of breast cancer risk over age, For panels A, B, C, and D, the Kaplan-Meier curves for each of the four VNTRs with 0.05 < FDR < 0.25 are shown. Panel A is VNTR558420, Panel B is VNTR 945060, Panel C is VNTR735300, and panel D is VNTR549198. The black line is the reference genotype and the red is the risk genotype. For each of the VNTRs, there were significantly different risks by genotype (log-rank p value < 0.05) with earlier ages of developing breast cancer among participants carrying the risk genotypes.
