## Supplemental Figure 3 for "Variable Number Tandem Repeats (VNTRs) as modifiers of breast cancer risk in carriers of *BRCA1* 185delAG"

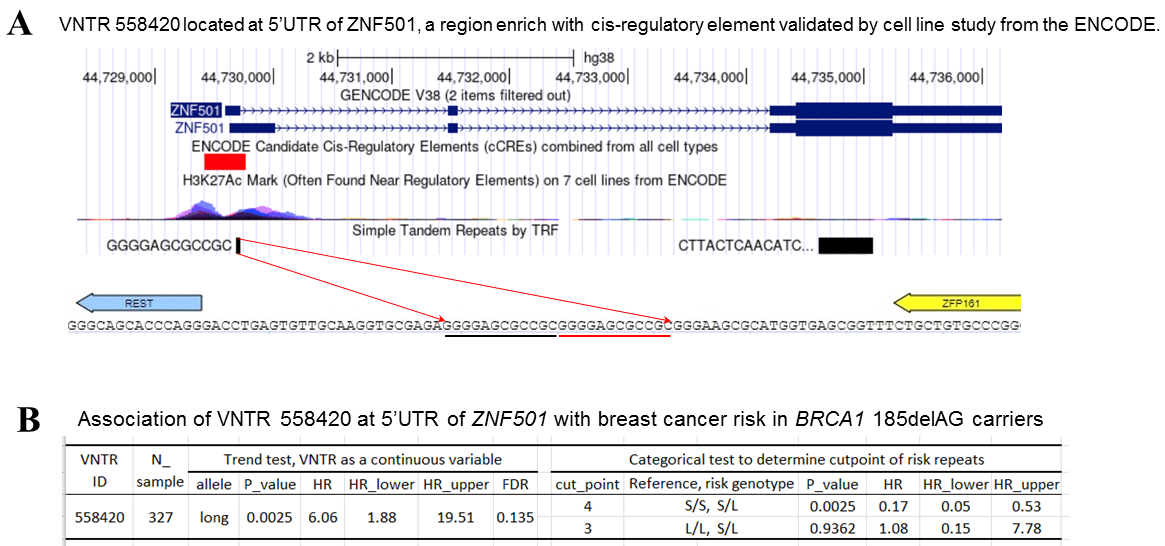

**Supplemental Figure 3.** VNTR 558420 located at the 5’UTR of ZNF501, a region enriched with cis-regulatory elements validated by cell line studies from ENCODE**.**
