## Supplementary figures and images for "Variable Number Tandem Repeats (VNTRs) as modifiers of breast cancer risk in carriers of *BRCA1* 185delAG"

### Supplemental Figure 4

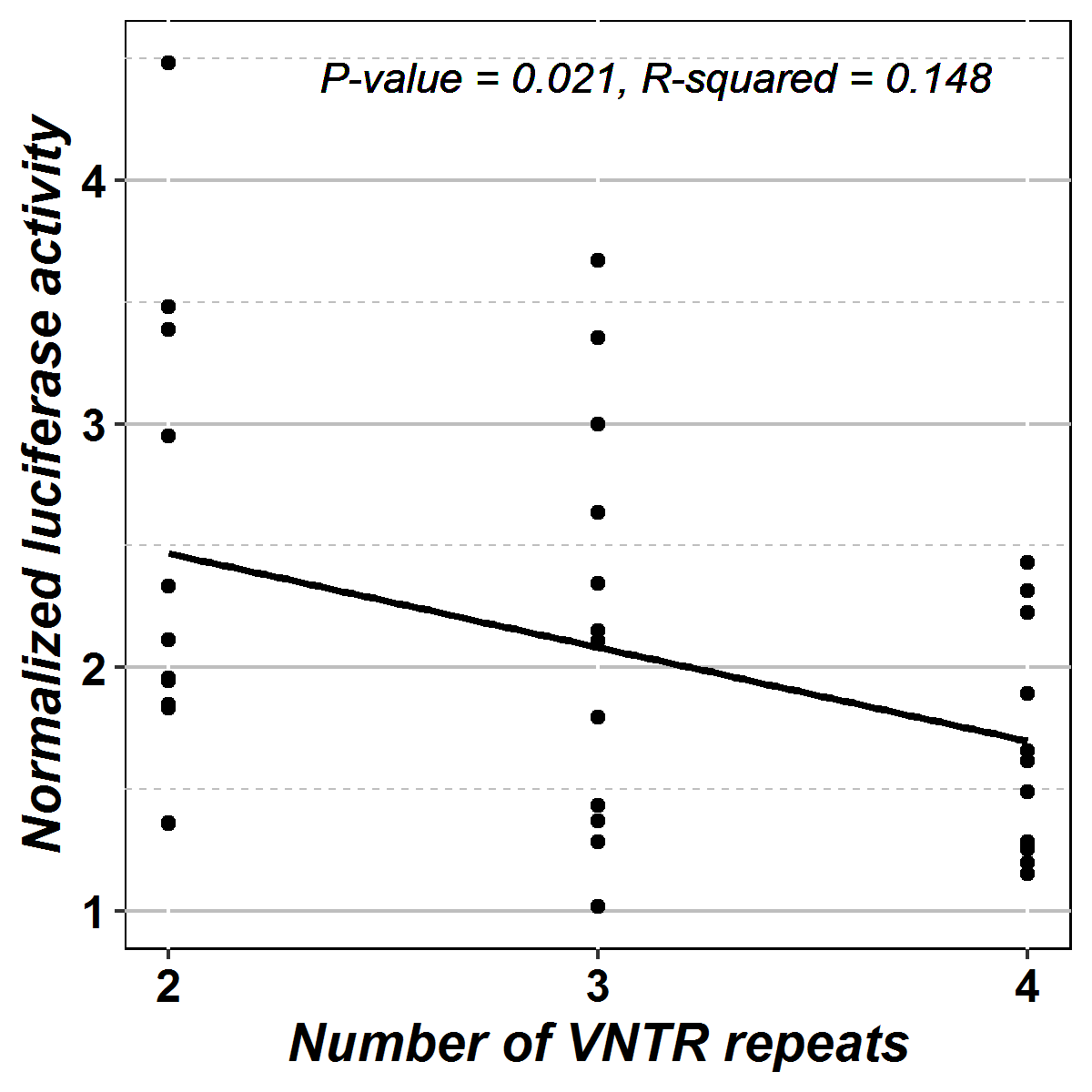

**Supplemental Figure 4.** A linear trend test of luciferase activity and number of repeats for VNTR558420.
