## Supplemental Figure 5 for "Variable Number Tandem Repeats (VNTRs) as modifiers of breast cancer risk in carriers of *BRCA1* 185delAG"

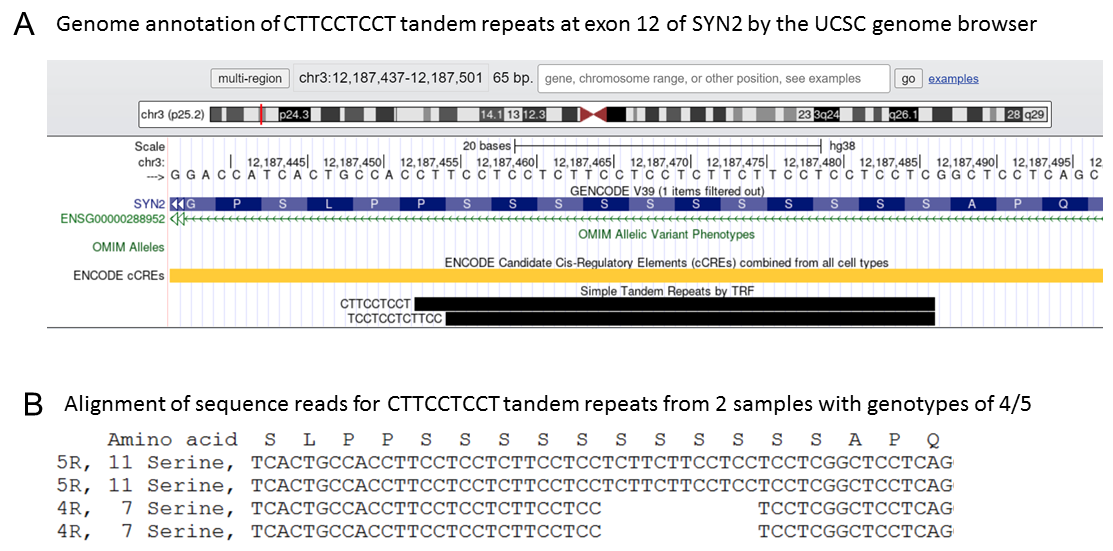

**Supplemental Figure 5:** Poly-Serine expansion resulted from by CTTCCTCCT tandem repeats at Exon 12 of SYN2**.** The Tandem Repeat Finder (TRF) program identified the repeat motif is CTTCCTCCT (panel A), the actual repeat motif is TCN (TCC, TCG, or TCC) (panel B).
