## Supplemental Figure 6 for "Variable Number Tandem Repeats (VNTRs) as modifiers of breast cancer risk in carriers of *BRCA1* 185delAG"

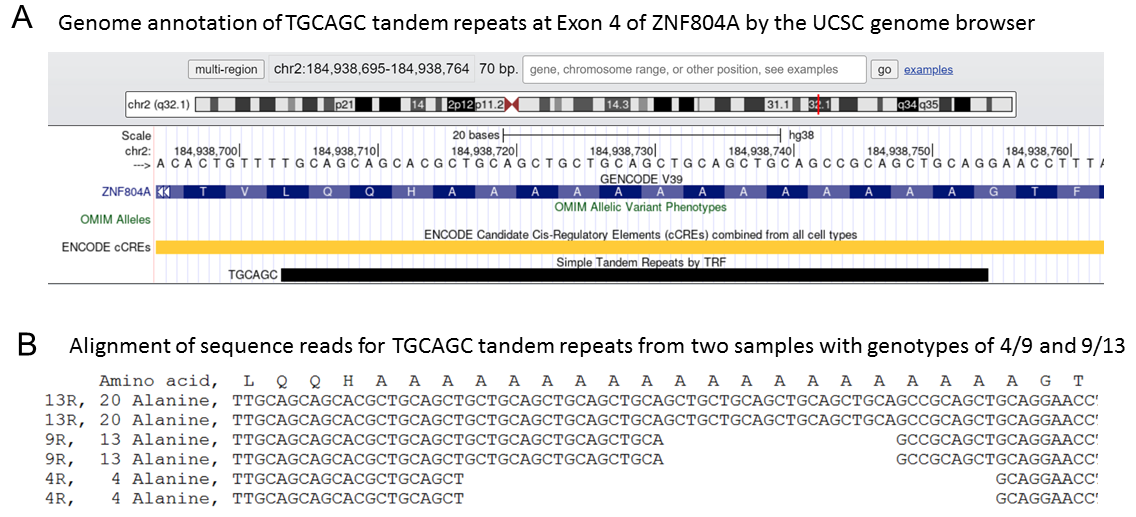

**Supplemental Figure 6**: Poly-Alanine expansion resulted from TGCAGC tandem repeats at Exon 4 of ZNF804A. The Tandem Repeat Finder (TRF) program identified the repeat motif is TGCAGC (panel A), the actual repeat motif is GCN (GCT, GCA, or GCC) (panel B).
