## Supplemental Table 1 for "Variable Number Tandem Repeats (VNTRs) as modifiers of breast cancer risk in carriers of *BRCA1* 185delAG"

Supplemental Table 1. Annotation of eight VNTRs (from Table 3) associated with risk of developing breast cancer.

| ID | Chr(hg38) | Start | End | Gene | Location**^a^** | Motif | Motif length (bp) |
| --- | --- | --- | --- | --- | --- | --- | --- |
| 253688 | 14 | 61654924 | 61655060 | FLJ22447 | 3' DS | GAAT | 4 |
| 357331 | 17 | 77880699 | 77880839 | LINC01973 | Exon 2 | GAGGCAGG | 8 |
| 472060 | 2 | 184938703 | 184938855 | ZNF804A | Exon 4 | TGCAGC | 6 |
| 412033 | 19 | 53600134 | 53600272 | LOC284379 | Exon 4 | AACA | 4 |
| 558420 | 3 | 44729686 | 44729823 | ZNF501 | 5'UTR | GGGGAGCGCCGC | 12 |
| 735300 | 6 | 42868874 | 42868999 | BICRAL | 3' DS | ATTTT | 5 |
| 945060 | X | 72238943 | 72239099 | ERCC6L | 5'UTR | GGAGCTT | 7 |
| 549198 | 3 | 12187452 | 12187596 | SYN2 | Exon12 | CTTCCTCCT | 9 |

**a**: 3’DS = 3’ downstream of a gene.
